# Split Selectable Markers

**DOI:** 10.1101/452979

**Authors:** Nathaniel Jillette, Menghan Du, Albert Cheng

**Affiliations:** The Jackson Laboratory for Genomic Medicine, Farmington, CT 06032, USA; Department of Genetics and Genome Sciences, University of Connecticut Health Center, Farmington, CT 06030, USA; Institute for Systems Genomics, University of Connecticut Health Center, Farmington, CT 06030, USA

## Abstract

Selectable markers are widely adopted in transgenesis and genome editing for selecting engineered cells with desired genotype but are limited in choices. We present here split selectable markers each allowing for selection of multiple “unlinked” transgenes in the context of lentivirus-mediated transgenesis as well as CRISPR/Cas-mediated biallelic knock-ins. Future development of split selectable markers may support enrichment or selection of “hyper-engineered” cells containing tens of transgenes or genetic modifications.

## Main

Selectable markers, such as antibiotic resistance or fluorescent protein genes, are often used in genetic engineering to isolate cells with desired genotypes ^1^. However, there are a limited number of well-characterized antibiotic resistance genes for use in eukaryotic cells and a limited number of fluorescent proteins whose spectra can be unambiguously differentiated by commonly used equipment. Researchers often run into the problem of not having enough choices of selectable markers if they are to incorporate multiple transgenes into a cell. On the other hand, selection with multiple antibiotics at the same time is often harsh to cells. “Selectable marker recycling” may provide a work-around, however, requiring multiple rounds of transgenesis, selection and removal of selection markers ^2^. To allow multiple transgenes to be selected by one selection scheme, we have created split antibiotics resistance and fluorescent protein genes wherein a gene encoding an antibiotic resistance or fluorescent protein is split into two or more segments fused to inteins (“markertrons”) that can be rejoined by protein trans-splicing ^3^ (Fig 1a). Each markertron is inserted onto a transgenic vector carrying a specific transgene. Delivery of transgenic vectors containing a set of markertrons yields cells harboring a subset or a complete set of the marketrons. Only cells containing a complete set of markertrons produce a fully reconstituted marker protein via protein splicing and thus passes through selection while cells with partial sets of markertrons are eliminated, achieving co-selection of cells containing all intended transgenes.

**Figure 1.**
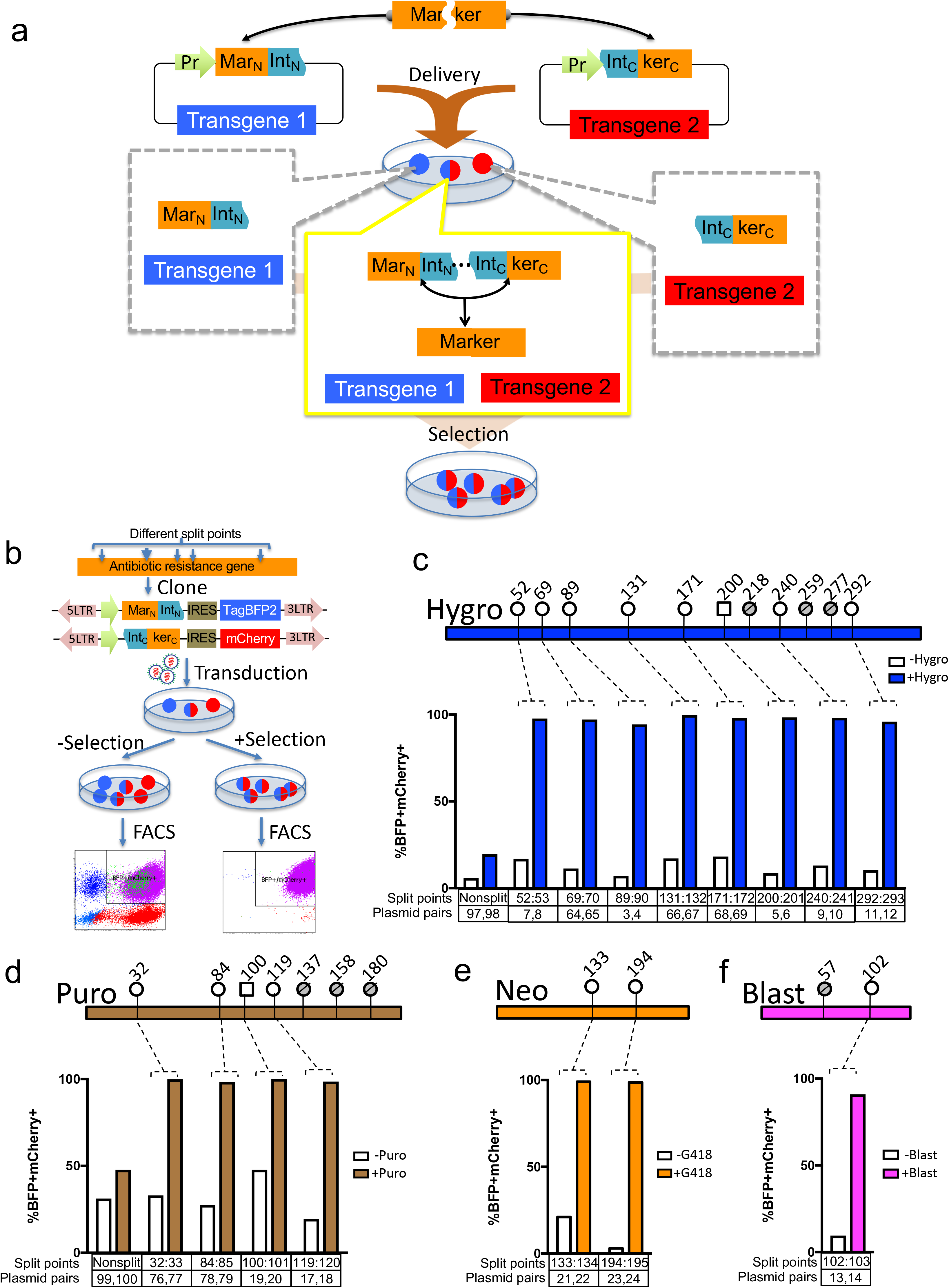
Split selectable marker for antibiotic co-selection of two separate transgenic vectors. (a) The coding sequence of a selectable marker is split into an N-terminal fragment (Mar_N_) and a C-terminal fragment (ker_C_) and separately cloned upstream of an N-terminal fragment of a split intein (Int_N_) and downstream of a C-terminal fragment of the split intein (Int_C_), respectively, on two different vectors each carrying a different transgene. These vectors are delivered to cells yielding sub-populations of cells containing either one of the vectors or both of the vectors. Only cells with both vectors expressing the two intein-split selectable marker fragments (“markertrons”) undergo protein trans-splicing to reconstitute a full-length selectable marker, allowing specific selection and enrichment of the double transgenic cells. (b) To screen for split points compatible for inteins for an antibiotic resistance gene, we identified potential split points according to the junctional requirement for the type of intein tested, then cloned the corresponding N-terminal and C-terminal fragments to the split intein scaffolds on lentiviral vectors equipped with TagBFP2 or mCherry fluorescent proteins, which serve as our test transgenes to evaluate selection efficiency. These are delivered into cells via lentiviral transduction. The cells were then split into replicate plates, one subjected to antibiotic selection while the other maintained in non-selective media. Following antibiotic selection, the replicate cultures were analyzed by flow cytometry. (c) 2-markertron hygromycin (Hygro) <u>int</u>ein-split <u>res</u>istance (Intres) genes. Top schematics shows the split points tested for hygromycin resistance gene. The last residue of the N-terminal fragment is indicated on top of the lollipops. Circle lollipops represent split points using *Npu*DnaE intein while square lollipops represent those using *Ssp*DnaB intein. Crossed-out and shaded lollipops indicate split pairs that failed to endow cells with hygromycin resistance. The column plot below shows the percentage of double-transgenic cells (BFP+ mCherry+) in the non-selected (white columns) and the selected cultures (blue columns) analyzed by flow cytometry. (d) 2-markertron puromycin (Puro) Intres genes. Top schematics show split points tested for puromycin resistance genes while bottom column plots show percentage of double transgenic cells in the non-selected (white columns) and the selected cultures (brown columns). (e) 2-markertron neomycin (Neo) resistance genes. Top schematics show split points tested for neomycin resistance genes while bottom column plots show percentage of double transgenic cells in the non-selected (white columns) and the selected cultures (orange columns). (f) 2-markertron blasticidin (Blast) Intres genes. Top schematics show split points tested for blasticidin resistance gene while bottom column plot shows percentages of double transgenic cells in the non-selected (white columns) and the selected (cyan columns) cultures.

We started out with engineering 2-markertron <u>int</u>ein-split <u>res</u>istance (Intres) genes for double transgenesis. Since flanking residues and local protein folding can affect efficiency of intein-mediated trans-splicing, we set out to identify split points in each of the four commonly used antibiotic resistance genes compatible with two well-characterized split inteins derived from *Npu*DnaE ^4,5^ and *Ssp*DnaB ^6^. To facilitate assessment of the effectiveness of double transgenic selection, we cloned markertrons onto lentiviral vectors expressing TagBFP2 or mCherry fluorescent proteins as test transgenes (Fig 1b). Viral preparations were transduced into U2OS cells, which were then split into replicate plates with non-selective or selective media. Following appropriate passages for antibiotics selection, the two cell cultures were analyzed by flow cytometry. For hygromycin (Hygro) resistance gene, one “native” *Ssp*DnaB split point (G200:S201) with flanking residues “GS” and one “native” *Npu*DnaE split point (Y89:C90) with “YC” residues were tested. Both enabled successful selection when both N- and C-markertrons were transduced yielding >99% BFP+ mCherry+ double transgenic cells in selected cultures compared to <10% double-positive cells in non-selected culture (Fig 1c; Plasmid pairs 3,4 and 5,6). Cells transduced with either of the two markertrons did not survive hygromycin selection. In contrast, double transgenesis with conventional full-length non-split hygromycin vectors only allow for ~20% enrichment of BFP+ mCherry+ cells (Plasmid pairs 97,98). We screened three addition potential split points (52S:53C),(240A:241C), and (292R:293C) for *Npu*DnaE with the obligatory cysteine residue on the C-extein junction and a residue on the N-extein junction that supported substantial trans-splicing activities in a previous report ^7^. We also incorporated six additional *Npu*DnaE split points by inserting an “artificial” cysteine on the C-extein junction to support splicing at ectopic sites yielding additional split points. In total, eight out of eleven split points tested supported hygromycin selection (Fig 1c). Similarly, for puromycin (Puro) (Fig 1d), neomycin (Neo) (Fig 1e) and blasticidin (Blast) (Fig 1f) resistance genes, we identified four, two, and one functional Intres pair(s), respectively. In all of these cases, cells transduced with either markertrons did not survive selection, while cells transduced with both yielded >95% double transgenic cells in selective cultures compared to <50% in non-selective cultures with the exception of Blasticidin(102) Intres, achieving lower but still significant enrichment of 91% double transgenic cells (Fig 1c~f). Details of the split points of Intres genes and plasmids are presented in Supplementary fig 1~4 and Supplementary table 1. To facilitate adoption of Intres markers, we created Gateway-compatible lentiviral vectors for convenient restriction-ligation-independent LR clonase recombination of transgenes ^8^ (Supplementary fig 5). We tested the functionality of these vectors by recombining TagBPF2 and mCherry, respectively to the N- and C-Intres vectors and found robust selection of double transgenic cells (Supplementary fig 5b). One potential utility of Intres vectors is to install different fluorescent markers in cells to label different cellular compartments. To explore such utility, we cloned in NLS-GFP and LifeAct-mScarlet ^9^, which label nucleus and F-actin, respectively, by Gateway recombination to conventional full length (FL) non-split hygromycin selectable vectors or 2-markertron hygromycin Intres vectors and transduced cells with either sets of plasmids, followed by antibiotic selection (Supplementary fig 5c). The sample transduced with non-split selectable plasmids contained both singly and doubly labelled cells, while cell transduced with Intres plasmids were all doubly labelled (Supplementary fig 5c).

To test whether split fluorescent markers can be used for transgene selection, we screened for *Npu*DnaE split points for mScarlet fluorescent protein (Supplementary fig 6 and 7a) and identified four split points allowing for >96% enrichment of double transgenic cells and three other split points enabling >60% enrichment of double transgenic cells in mScarlet-gated population, compared to <20% double transgenic cells in non-gated population (Supplementary fig 7b).

With the split points identified for 2-markertron Intres genes, we set out to engineer higher degree split markers. We tested combinations of splits points to partition a marker gene into three or more markertrons to allow for co-selection of more than two “unlinked” transgenes with one antibiotic (Fig 2a and b). To identify pairs of split points that would allow such “Intres chain”, we cloned 3-split markertrons into three lentiviral vectors each carrying one of three fluorescent transgenes TagBFP2, EGFP, or mCherry, that will allow us to assess effectiveness of selection by flow cytometry (Fig 2c). Since hygromycin resistance gene is the longest and provides the most split points for testing, we focused on engineering 3-markertron hygromycin Intres. We tested two 3-markertron hygromycin Intres using two intervening *Npu*DnaE inteins, two using *Npu*DnaE for the first intein and *Ssp*DnaB for the second intein, as well as two using *Ssp*DnaB for the first intein and *Npu*DnaE for the second intein (Fig 2d). Five of these six 3-markertron hygromycin Intres enabled >97% and with the remaining one enabling 80% triple transgenic selection in hygromycin-selected cultures compared to <15% triple transgenic cells in non-selected cultures. Samples with leave-one-out transduction did not yield any viable cells after hygromycin selection while cells transduced with non-split hygromycin vectors yielded only 7% triple transgenic cells after selection. To facilitate the use of 3-markertron Intres, we created Gateway compatible lentiviral vectors with these markers (Supplementary fig 8a). Three sets of these vectors were each tested by recombining TagBFP (as transgene 1), EGFP (as transgene 2) and mCherry (as transgene 3) into the N-, M-, and C-Intres Gateway destination vectors and used to transduce U2OS cells, which were then split and cultured in hygromycin selection or non-selective media (Supplementary fig 8b). Two weeks after selection, cells were analyzed by flow cytometry. All three sets of 3-markertron hygromycin Intres plasmids support triple transgenic cell selection of >99% compared to <25% in the non-selected cultures (Supplementary fig 8c).

**Figure 2.**
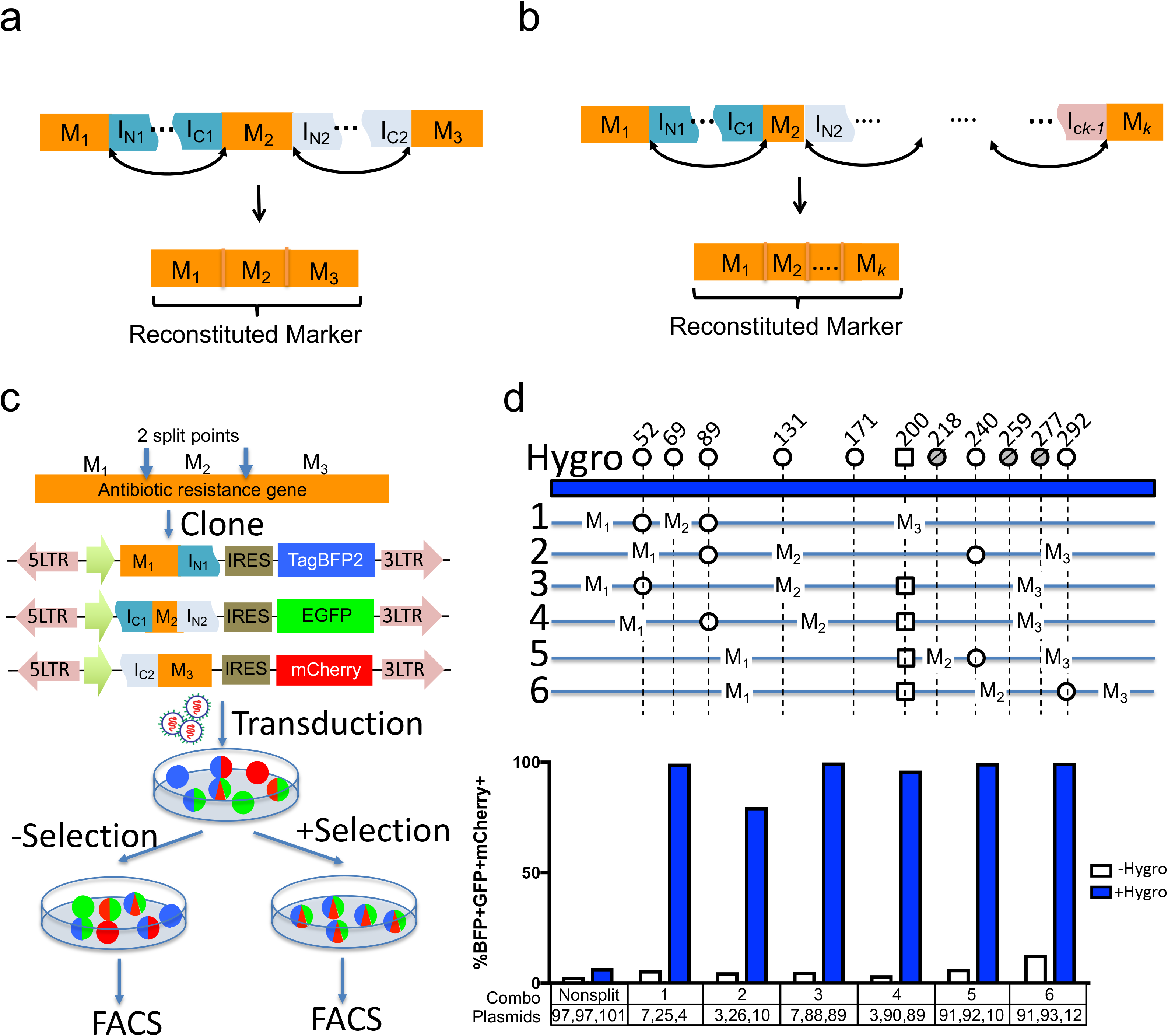
Multi-split selectable markers for co-selection of three or more transgenic vectors. (a) A selectable marker is partitioned into three fragments (M_1_, M_2_ and M_3_). The first marker fragment (M_1_) is fused upstream of the N-terminal fragment of the first split intein (I_N1_). The second marker fragment (M_2_) is fused downstream of the C-terminal fragment of the first split intein (I_C1_) and upstream of the N-terminal fragment of the second split intein (I_N2_). The third marker fragment (M_3_) is fused downstream of the C-terminal fragment of the second split intein (I_C2_). The first split intein catalyzes the joining of M_1_ to M_2_ while the second split intein catalyzes the joining of M_2_ to M_3_, effectively reconstituting the full selectable marker. (b) A design of a *k*-split selectable marker via an “intein chain” mechanism. Similar to the 3-split scenario, the selectable marker is partitioned into *k* fragments, and are reconstituted through protein trans-splicing mediated by intervening split inteins. (c) Split points identified from 2-split selectable markers were used in combination to produce 3-split selectable markers. The corresponding fragments were cloned into lentiviral vectors to result in the 3-split selectable marker structure and a reporter fluorescent transgene per vector. Cells were then transduced with viruses prepared from these vectors, split into selective or non-selective media. After appropriate selection period, the cultures were analyzed by flow cytometry. (d) 3-markertron hygromycin (Hygro) Intres. Top schematics shows the split points tested for hygromycin resistance gene, with residue numbers of the last amino acid of the N-terminal fragments indicated above circle or square lollipops, representing *Npu*DnaE and *Ssp*DnaB inteins, respectively. Six 3-markertron hygromycin Intreses were tested, each indicated with a numbered line with circle or square indicating the two split points used for each case. Column plot below shows the percentage of triple transgenic (BFP+ GFP+ mCherry+) cells from the non-selective (white columns) and selective (blue columns) cultures for the 3-markertron hygromycin Intres indicated by the numbers below.

A further potential application of split selectable markers is to facilitate genome engineering and editing via the CRISPR/Cas system ^10^. Although gene knockout based on NHEJ-mediated insertions/deletions (indels) occur at high frequency, precise editing and knock-in based on homology directed repair (HDR) using exogenous repair templates are inefficient ^11^. We tested whether split selectable markers can be used to select for cells with CRISPR-mediated biallelic knock-in at the *AAVS1* locus ^12^. We constructed targeting constructs with homology arms flanking the target site, and splice acceptor-2A peptide to trap the markertrons within intron one of the host gene *PPP1R12C*. However, we did not obtain any live cells after CRISPR/Cas knock-in experiments in HEK293T cells using these targeting constructs and two weeks of antibiotic selection (Data not shown). We suspected that the endogenous promoter of the host gene *PPP1R12C* might not drive sufficient expression of markertrons to reconstitute enough antibiotic resistance protein to counteract actions of the antibiotics. We thus tested an alternative strategy to express Intres markertrons by TetO promoter whose activity can be titrated by doxycycline (dox) concentration. To allow comparison of Intres-mediated biallelic selection versus full-length (FL) non-split selectable markers, we implemented several different targeting construct designs. First, we drive expression of a full-length (FL) resistance gene (e.g., Hygro) together with rtTA under a constitutive EF1a promoter and a separate test Intres (e.g., Blast Intres) under a dox-inducible TetO promoter (Supplementary fig 9b, Plasmids 109 and 110). This allows comparison of full length and split selectable markers within the same constructs. To allow fair comparison of full length versus split markers driven by the same TetO promoter, we constructed two similar plasmids 107 and 108 (cf. Plasmids 109 and 110), wherein full-length antibiotic resistance gene (Blast) is placed downstream of the TetO promoter. To enable single-cell quantification of biallelic targeting and to demonstrate the feasibility of incorporating two transgenes into two *AAVS1* alleles, we appended EGFP and mScarlet fluorescent genes downstream of the test split or non-split markers via self-cleaving 2A peptide. Similarly, to test Hygro Intres, we swapped the EF1a and TetO-driven markers so that FL Hygro or Hygro Intres were placed downstream of TetO and FL Blast downstream of EF1a (Supplementary fig 9c and d; Plasmids 111~114). We co-transfected pX330-AAVS1 (Plasmid 106) containing Cas9 and sgRNA targeting *AAVS1*, and the different pairs of targeting constructs (TC) to HEK293T cells, split into triplicate doxycycline-containing media without antibiotics, with blasticidin, or with hygromycin at the subsequent passages. Two weeks after selection, we analyzed the cultures for biallelic targeting by flow cytometric measurement of GFP and RFP fluorescence (Supplementary fig 9e). As expected, non-selected cultures harbored small fraction (<1%) of biallelic knock-in GFP+/RFP+ cells (Supplementary Fig 9e; Selection = None). Selection of antibiotics where corresponding FL antibiotic resistance genes were present on targeting constructs yielded < 30% biallelic knock-in cells (Supplementary fig 9e; Blast: TC a,c,d; Hygro: TC a,b,c). In contrast, selection of antibiotics where corresponding Intres are present on the targeting constructs yielded 75% (Supplementary fig 9e; Blast Intres: TC b) and 88% (Supplementary fig 9e; Hygro Intres: TC d) biallelic knock-in cells. Selection for an additional two weeks allowed split Blast and Hygro TCs to achieve 96.5% and 97.0% biallelic knock-in, respectively (Supplementary fig 9f and g). We next tested bilallelic engineering in KOLF2-C1 human induced pluripotent stem cells (hiPSCs), which are karyotypically normal with a stable diploid genome ^13^ (Fig 3). The full-length non-split Blast targeting constructs (Fig 3a) and 2-split Blast Intres targeting constructs (Fig 3b) were tested for selection of biallelically modified clones. Purified Cas9 proteins were complexed with synthetic sgRNA to form Cas9 ribonucleoprotein (RNP) and co-nucleofected with the targeting constructs into KOLF2-C1, followed by dox-induction and antibiotic selection. Surviving colonies were picked into separate wells for establishing single cell clones. Genotyping PCR revealed that targeting using non-split Blast resistance gene generated only 8% biallelic clones, while targeting using Blasticidin Intres yielded exclusively (100%) biallelically modified clones (Fig 3c and d), showing both fluorescent signals (Fig 3e) indicative of the targeting by each targeting construct at the two alleles of AAVS in these hiPSCs.

**Figure 3.**
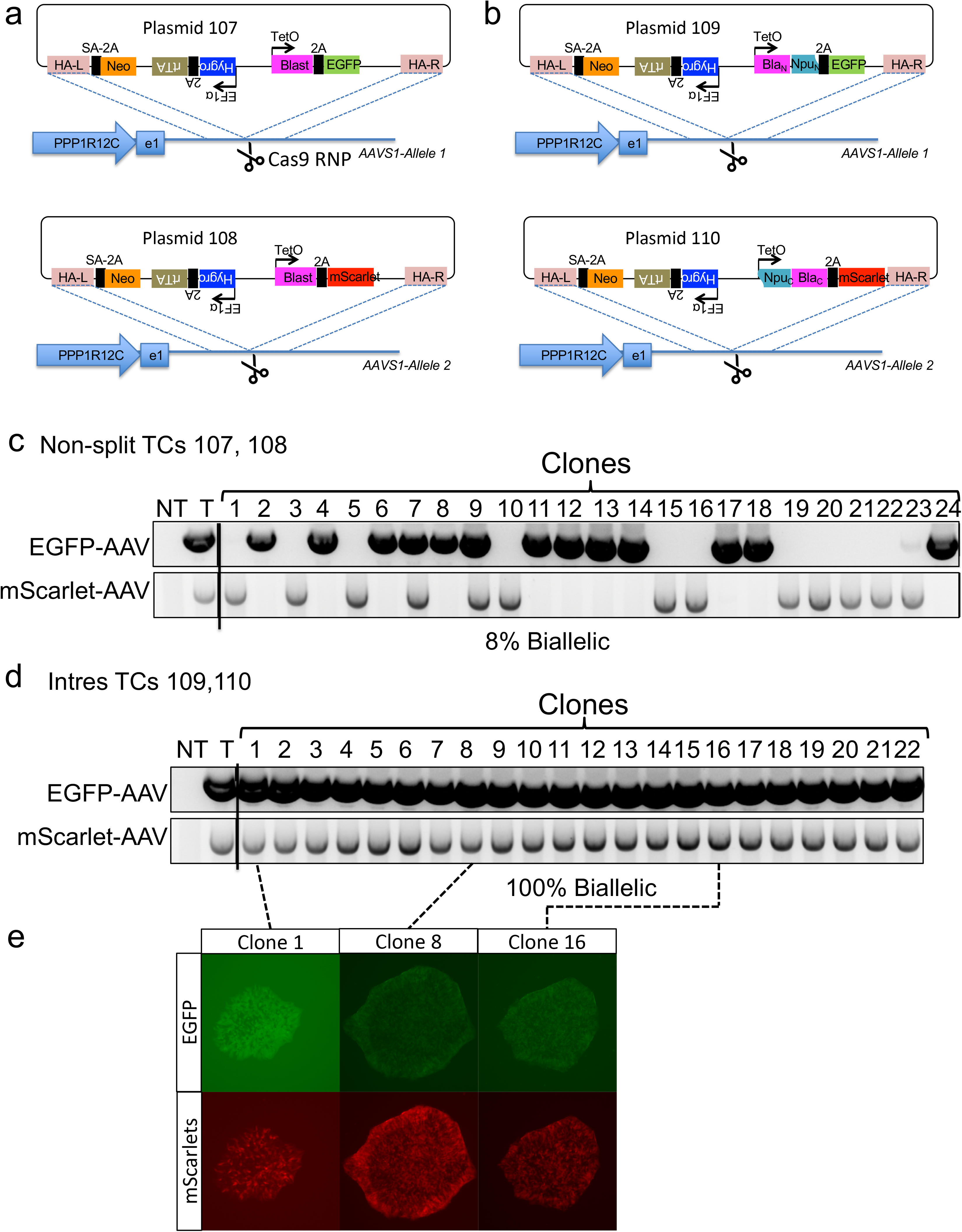
Intres for selection of biallelic CRISPR knock-in in human induced pluripotent stem cells. (a) Plasmids 107 and 108 contains AAVS homology arms, and Dox-inducible full length (FL) Blasticidin (Blast) and EGFP (plasmid 107) or mScarlet (plasmid 108) separated by a self-cleaving 2A peptide sequence. (b) Plasmids 109 and 110 contains AAVS homology arms, and Dox-inducible N-split Blast markertron and EGFP (plasmid 109) or C-split Blast markertron and mScarlet (plasmid 110) separated by a self-cleaving 2A peptide sequence. Cas9 ribonucleoprotein was formed *in vitro* by complexing purified Cas9 proteins with synthetic sgRNA targeting AAVS locus, introduced to hiPSCs by nucleofection, subsequently subjected to dox-induction and antibiotic selection, and finally cloned by colony picking and analyzed for allelic modification. (c) Genotyping PCR for EGFP- and mScarlet-inserted AAVS alleles of single iPSC clones generated by CRISPR editing using non-split targeting constructs (TCs) 107 and 108. (d) Genotyping PCR for EGFP- and mScarlet-inserted AAVS alleles of single iPSC clones generated by CRISPR editing using Intres TCs 109 and 110. (e) Representative fluorescent microscopic images of hiPSC colonies from the indicated clones derived from the Intres CRISPR experiments.

In this study, we have engineered split antibiotic resistance and fluorescent protein genes that can allow selection for two or more “unlinked” transgenes. By inserting unnatural residues at selectable markers, we showed that novel high-efficiency split points could be utilized, expanding the positions available for engineering. We demonstrated that split selectable markers could be incorporated into lentiviral vectors or gene targeting constructs in CRISPR/Cas9 genome editing experiments for positive selection of cells with double transgenesis or biallelic knock-ins. By combining two or more splits points, we showed that 3- and 4-split markers could be generated to allow higher degree transgenic selection. Future development of even higher-degree split selectable markers may enable “hyper-engineering” of cells containing tens of transgenes or targeted knock-ins.

## Acknowledgment

Research reported in this publication was partially supported by the Jackson Laboratory internal grants, National Human Genome Research Institute (1R01HG009900) and National Cancer Institute (P30CA034196). KOLF2-C1 cells were a gift from Bill Skarnes and were derived from the HipSci consortium.

## Supplementary Information

**Supplementary figure 1.**
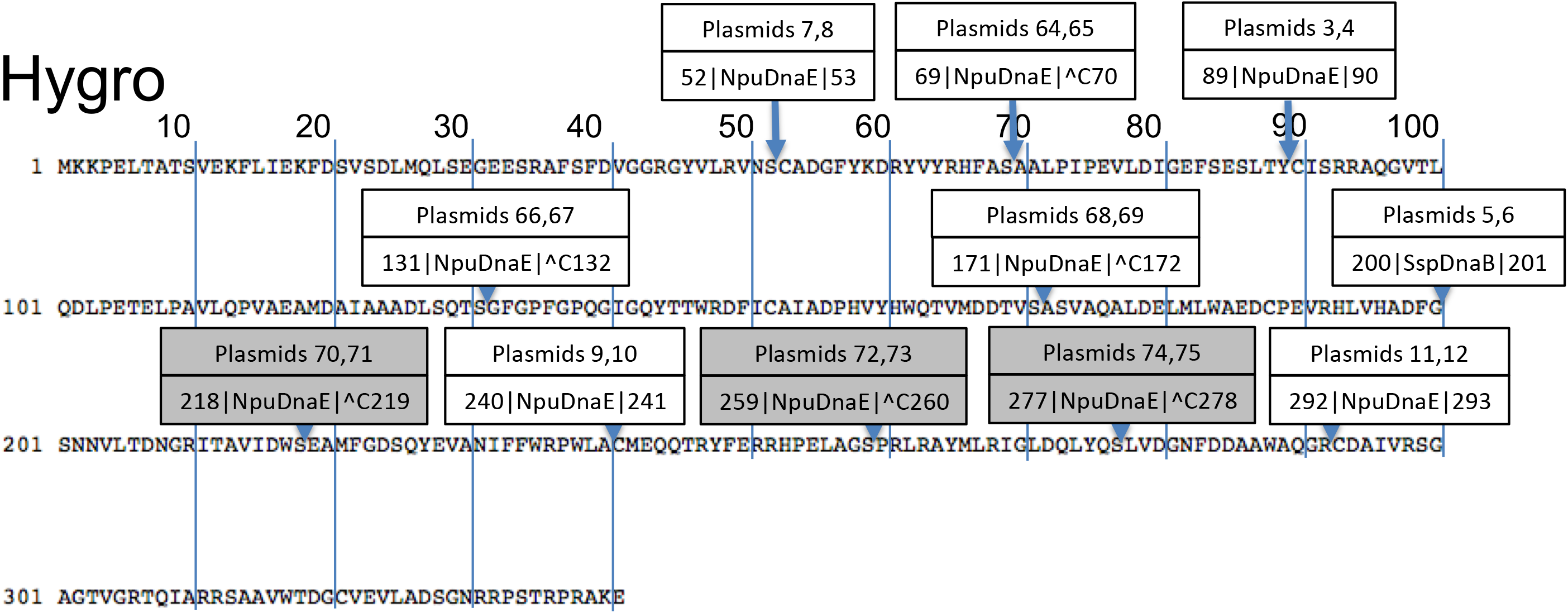
Split points for hygromycin resistance protein. Amino acid sequence of hygromycin resistance protein is presented with clouds labeling the split points characterized in this study. Within the label, the top row indicates the plasmid numbers corresponding to Supplementary Table 1. The bottom row indicates the residue number of the last amino acid in the N-terminal fragment, the species of the intein used, and the residue number of the first amino acid in the C-terminal fragment. “^C” indicates an insertion of a Cysteine.

**Supplementary figure 2.**
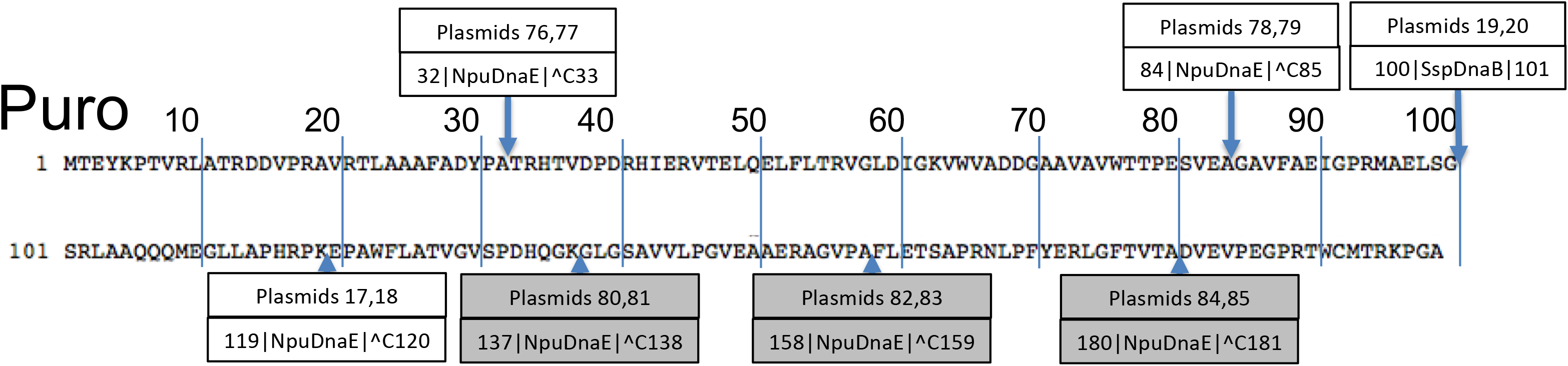
Split points for puromycin resistance protein. Amino acid sequence of puromycin resistance protein is presented with clouds labeling the split points characterized in this study. Within the label, the top row indicates the plasmid numbers corresponding to Supplementary Table 1. The bottom row indicates the residue number of the last amino acid in the N-terminal fragment, the species of the intein used, and the residue number of the first amino acid in the C-terminal fragment. “^C” indicates an insertion of a Cysteine.

**Supplementary figure 3.**
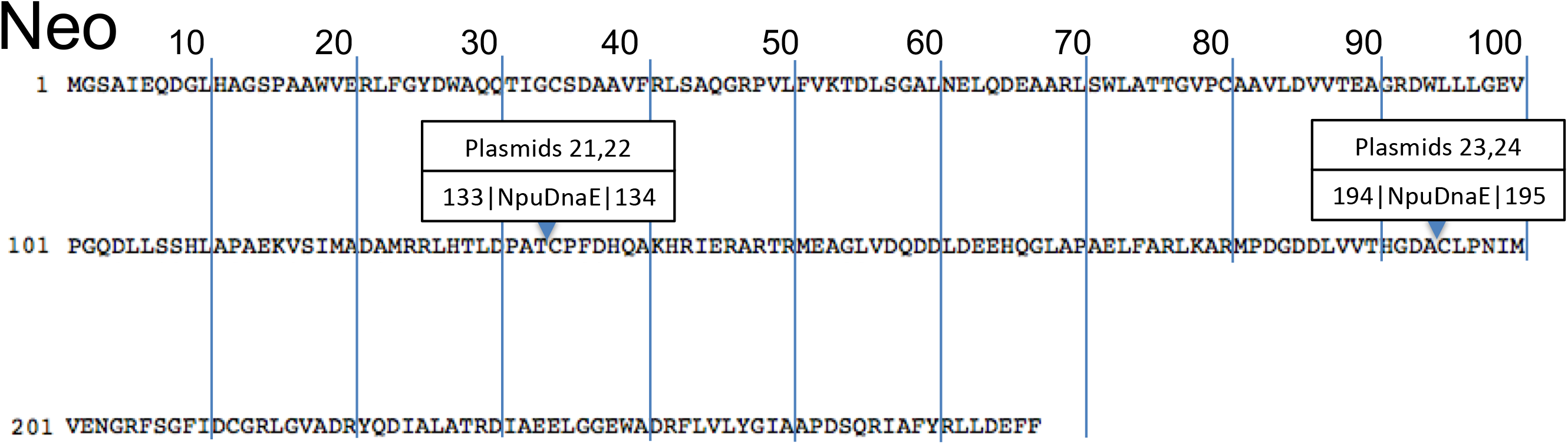
Split points for neomycin resistance protein. Amino acid sequence of neomycin resistance gene is presented with clouds labeling the split points characterized in this study. Within the label, the top row indicates the plasmid numbers corresponding to Supplementary Table 1. The bottom row indicates the residue number of the last amino acid in the N-terminal fragment, the species of the intein used, and the residue number of the first amino acid in the C-terminal fragment.

**Supplementary figure 4.**
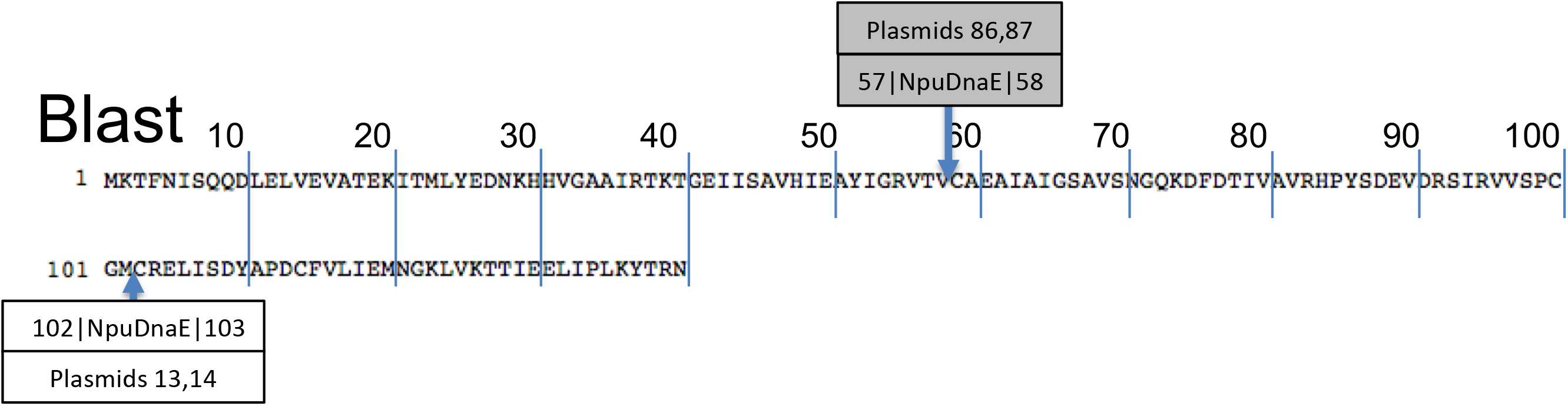
Split points for blasticidin resistance protein. Amino acid sequence of blasticidin resistance gene is presented with clouds labeling the split points characterized in this study. Within the label, the top row indicates the plasmid numbers corresponding to Supplementary Table 1. The bottom row indicates the residue number of the last amino acid in the N-terminal fragment, the species of the intein used, and the residue number of the first amino acid in the C-terminal fragment.

**Supplementary figure 5.**
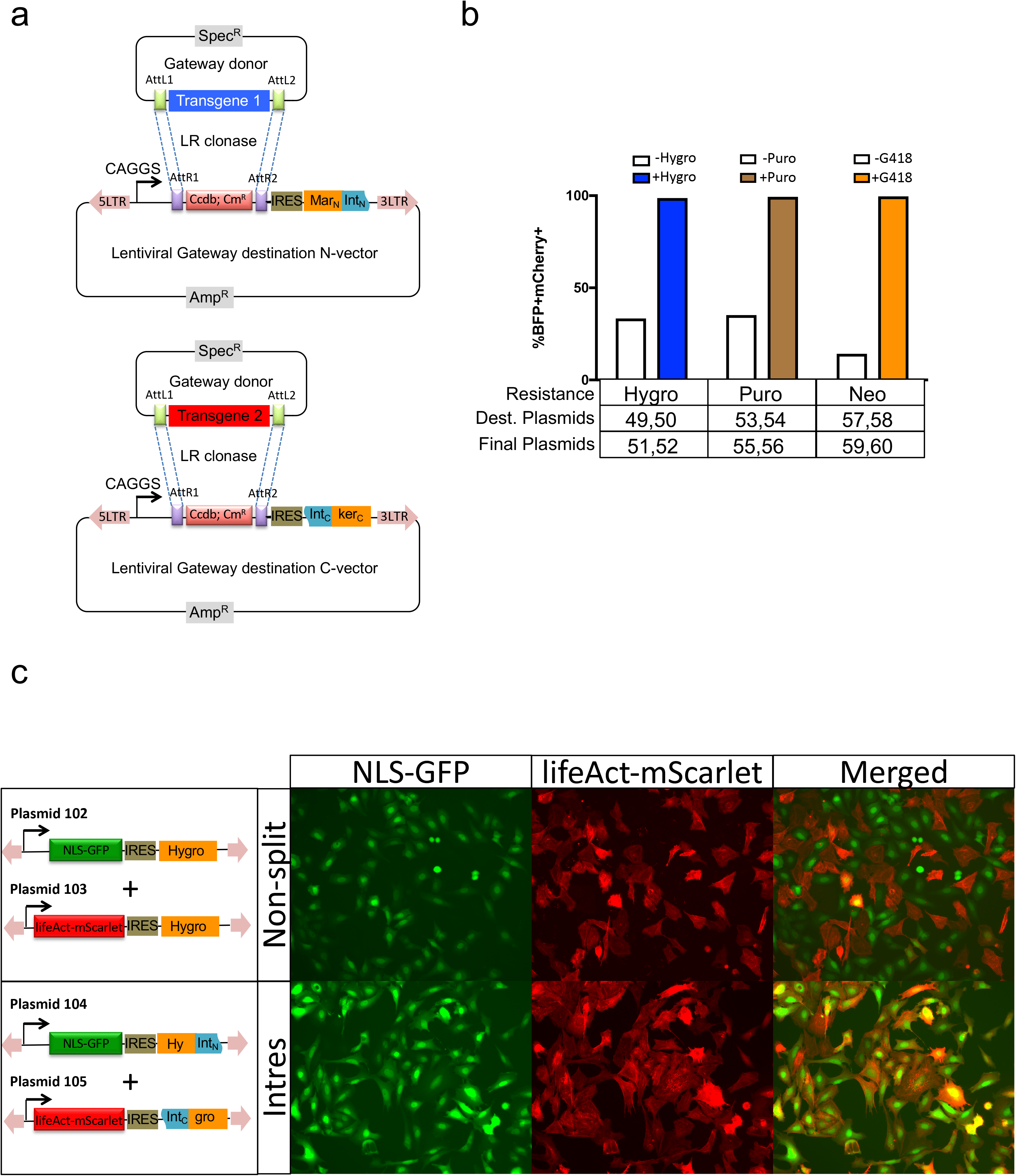
Gateway-compatible lentiviral destination vectors with 2-markertron Intres markers. (a) Gateway-compatiable lentiviral destination vector kits for each split Intres marker consists of an N-vector and C-vector. N-vector contains viral LTRs, CAGGS promoter, gateway destination cassette AttL, ccdB gene, chloramphenicol resistance gene that allows LR clonase-mediated recombination of Gateway donor vector carrying transgenes, followed by internal ribosomal entry site (IRES) that allows polycistronic expression of the N-markertron. Similarly, C-vector contains the C-markertron and allows recombination of another transgene. (b) TagBFP2 (as transgene 1) and mCherry (as transgene 2) were cloned into the 2-markertron Intres plasmids by Gateway recombination and delivered to cells by lentiviral transduction, followed by antibiotic selection and flow cytometry analysis. Column plot shows the percentage of BFP+mCherry+ double-positive cell in the selected culture from the 2-markertron hygromycin (Hygro, blue column), puromycin (Puro, brown column), and neomycin (Neo, orange column) experiments versus their corresponding non-selective cultures (white columns). (c) NLS-GFP (as transgene 1) that labels nucleus with GFP fluorescence and lifeAct-mScarlet (as transgene 2) that labels F-actin with mScarlet fluorescence were recombined into lentiviral vectors expressing full non-split hygromycin resistance gene or lentiviral vectors with 2-markertron hygromycin Intres genes and used to transduce U2OS cells to make dual-label cells. Representative fluorescence microscopic images show GFP, mScarlet and merged channels of cells after hygromycin selection for two weeks.

**Supplementary figure 6.**
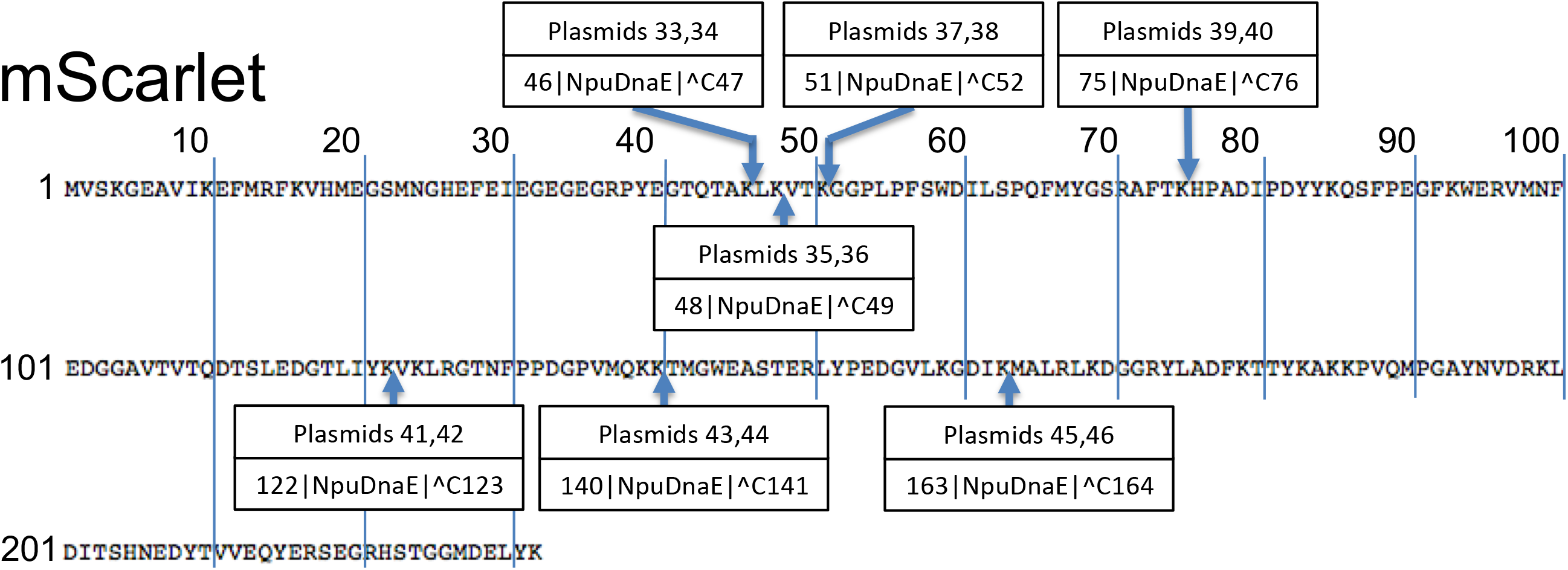
Split points for mScarlet fluorescent protein. Amino acid sequence of mScarlet gene is presented with clouds labeling the split points characterized in this study. Within the label, the top row indicates the plasmid numbers corresponding to Supplementary Table 1. The bottom row indicates the residue number of the last amino acid in the N-terminal fragment, the species of the intein used, and the residue number of the first amino acid in the C-terminal fragment. “^C” indicates an insertion of a Cysteine.

**Supplementary figure 7.**
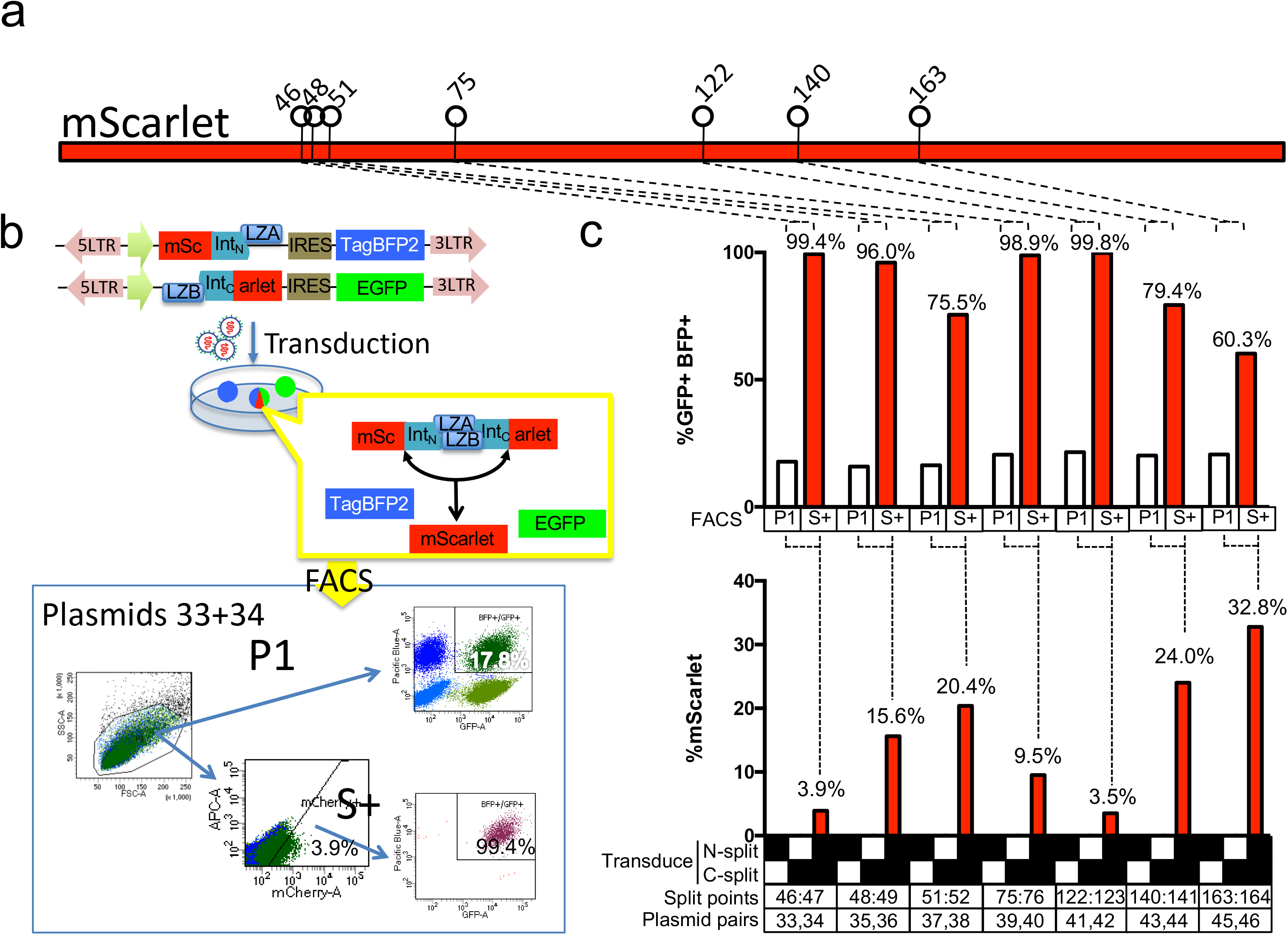
Split mScarlet for fluorescence-mediated co-selection of two separate transgenic vectors. (a) 2-markertron mScarlet proteins. Top schematics shows the split points tested for mScarlet. The last residue of the N-terminal fragment is indicated on top of the lollipops. (b) To screen for NpuDnaE intein-compatible split points for mScarlet, we identified potential split points according to the junctional requirement for *Npu*DnaE intein, then cloned the corresponding N-terminal and C-terminal fragments to the split inteins scaffolds on lentiviral vectors equipped with TagBFP2 or EGFP fluorescent proteins, which serve as our test transgenes to evaluate the selection efficiency. These are delivered into cells via lentiviral transduction. Cells with both lentiviruses contain the necessary protein splicing machinery and mScarlet fragments to reconstitute the full-length mScarlet fluorescent protein, as well as express both TagBFP2 and EGFP transgenes. Cells were subjected to FACS analysis. Boxed schematic shows an example of FACS analysis of the plasmid pair 33+34. P1 population was gated for forward scatter and side scatter for live single cells. From those, 17.8% of cells are double positive for TagBFP2 and EGFP transgenes. When the P1 cells were further gated for mScarlet-positive (mCherry channel), 99.4% of cells are double positive for TagBFP2 and EGFP. (c) The column plot below shows the percentage of mScarlet-positive cells of each of the indicated split points. The column plot above shows the percentage of TagBFP2+ EGFP+ cells among the P1 cells (white columns) and the mScarlet-positive subset of P1 cells (red columns).

**Supplementary figure 8.**
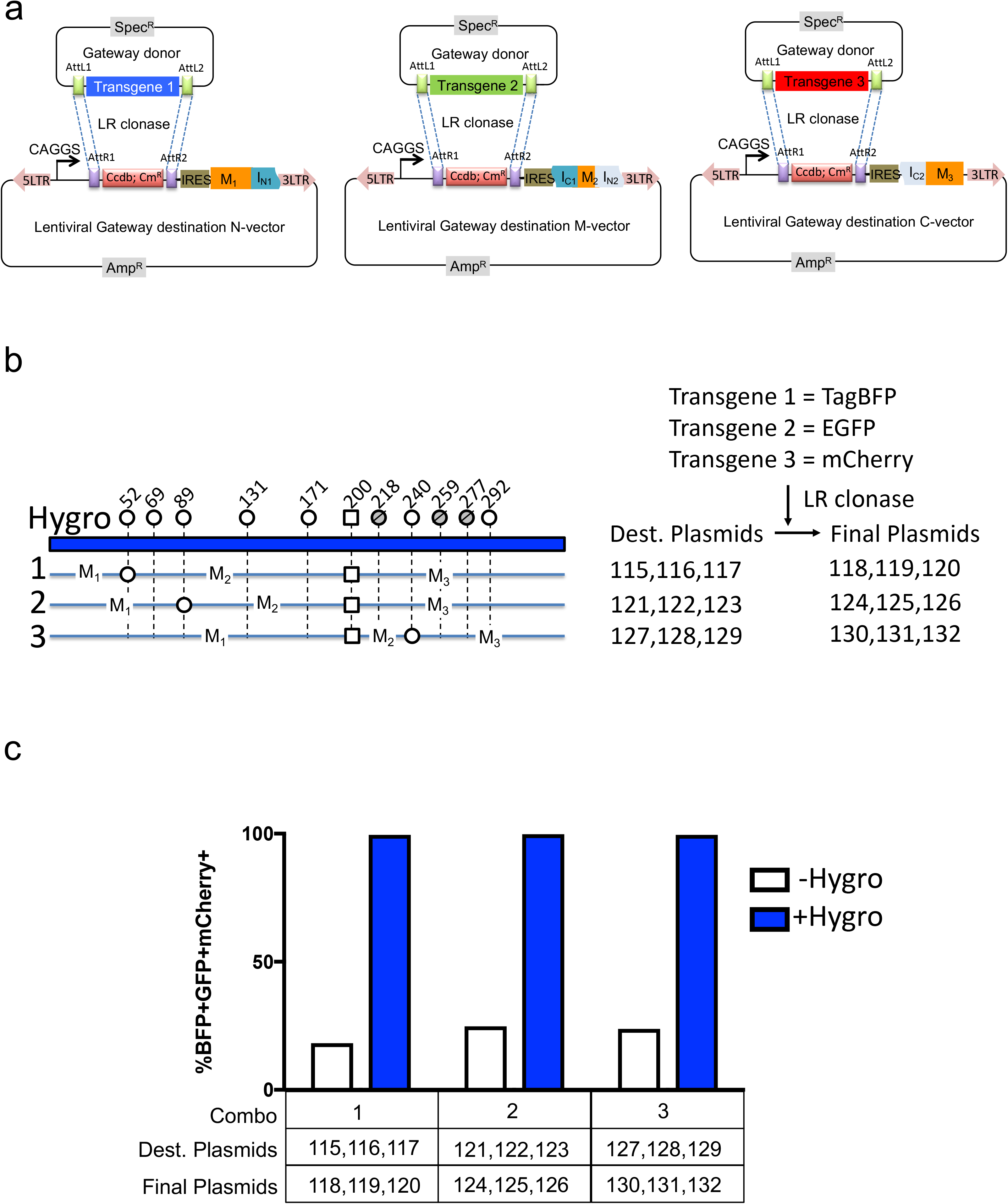
Gateway-compatible lentiviral destination vectors with 3-markertron hygromycin Intres genes. (a) Gateway-compatiable lentiviral destination vector with viral LTRs, CAGGS promoter, gateway destination cassette AttL, ccdB gene, chloramphenicol resistance gene that allows LR clonase-mediated recombination of Gateway donor vector carrying transgenes, followed by internal ribosomal entry site (IRES) that allows polycistronic expression of the each of the three 3-split hygromycin markertrons. (b) TagBFP2 (as transgene 1) and EGFP (as transgene 2) and mCherry (as transgene 3) were cloned into the 3-split Intres plasmids by Gateway recombination and delivered to cells by lentiviral transduction, followed by antibiotic selection and flow cytometry analysis. (c) Column plot shows the percentage of BFP+GFP+mCherry+ triple-positive cells in the hygromycin selected (blue columns) versus their corresponding non-selective cultures (white columns).

**Supplementary figure 9.**
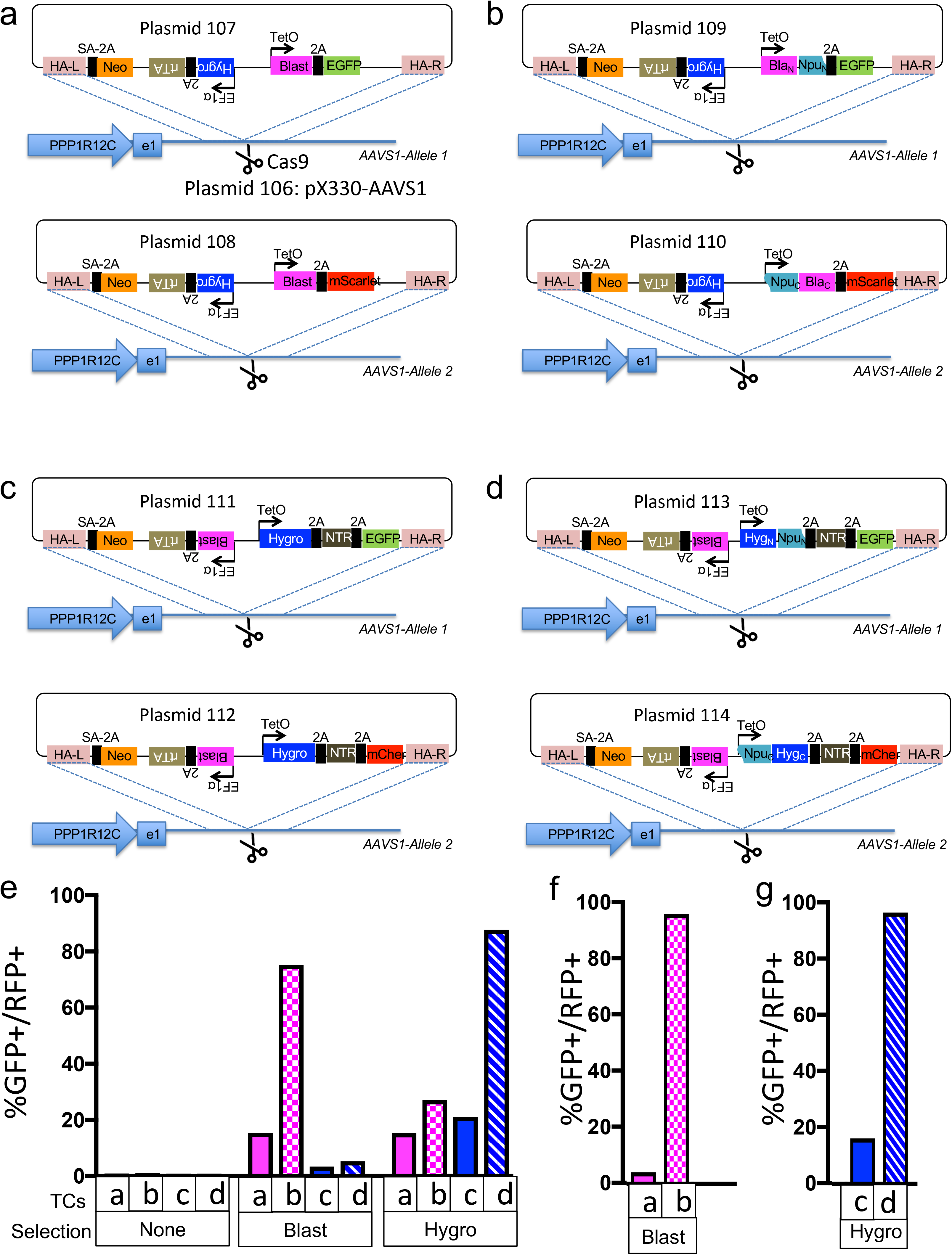
Intres markers allow enrichment of biallelic targeted cells from CRISPR/Cas-mediated knock-in experiments. Targeting construct pairs containing homology arms for AAVS1 safe harbor locus were designed to contain full length (FL) non-split or split Intres markers and tested for ability to enrich for biallelic targeted cells via antibiotic selection. (a) Plasmids 107 and 108 contains FL Neomycin (Neo) resistance gene driven by endogenous PPP1R12C promoter at the AAVS1 locus, FL Hygromycin (Hygro) gene and rtTA Dox-respsonsive transactivator driven by an EF1a promoter, as well as FL Blasticidin (Blast) expressed as well as EGFP (plasmid 107) and mScarlet (plasmid 108) from a dox-inducible TetO promoter. Plasmid 106 contains Cas9 and an sgRNA targeting the AAVS locus. 2A: self-cleaving 2A peptides. Plasmids 106, 107 and 108 were co-transfected into HEK293T cells, split, and passaged in dox-containing hygromycin, blasticidin or non-selective media for two weeks, and analyzed by flow cytometry to assay efficiency of biallelic targeting. (b) Plasmids 109 and 110 contain similar structure as Plasmids 107 and 108, but having split Blast Intres in place of the FL Blast. (c) Plasmids 111 and 112 contain an EF1a-driven FL Blast and TetO-driven FL Hygro, nitroreductase (NTR), fluorescent protein (EGFP or mCherry) separated by 2A peptides. (d) Plasmids 113 and 114 are similar to Plasmids 111 and 112 but with Hygro Intres in place of FL Hygro. (e) Flow cytometry analysis of cells transfected with Plasmid 106 (Cas9+AAVS-sgRNA) and the indicated targeting construct pairs, two weeks after culturing in dox-containing non-selective media (Selection: None), blasticidin selection media (Blast) and hygromycin selection media (Hygro). (f) Flow cytometry analysis of cells transfected with Plasmid 106 (Cas9+AAVS-sgRNA) and the indicated targeting construct pairs, four weeks after culturing in dox-containing non-selective media (Selection: None) or blasticidin selection media (Blast) (g) Flow cytometry analysis of cells transfected with Plasmid 106 (Cas9+AAVS-sgRNA) and the indicated targeting construct pairs, four weeks after culturing in dox-containing non-selective media (Selection: None) or hygromycin selection media (Hygro).

## Online Methods

### Cloning

To generate a test plasmid for each markertron, we first generated a Gateway donor plasmid containing its ORF and then recombined into lentiviral destination vector with TagBFP2 (Plasmid 94: pLX-DEST-IRES-TagBFP2), EGFP (Plasmid 95: pLX-DEST-IRES-EGFP), or mCherry (Plasmid 96: pLX-DEST-IRES-mCherry) reporters, which were derived from pLX302 (Gift from David Root; Addgene: http://www.addgene.org/25896/) by removing Puromycin resistance gene and inserting IRES-fluorescent genes downstream of the Gateway cassette. The markertron-ORF Gateway donor plasmids were generated either by a nested fusion PCR procedure to combine intein with the coding sequence of fragments of the selectable marker followed by insertion into the pCR8-GW-TOPO plasmid by sequence- and ligation-independent cloning (SLIC) ^1^, or PCR-amplifying the relevant fragment of the selectable marker followed by insertion into “scaffold” plasmids (Plasmids 27~32) containing the intein sequences by SLIC. DNA sequences encoding inteins were codon optimized for *Homo sapiens,* and synthesized as GBlock (IDT), with AC1947GB encoding *Npu*DnaE intein, AC1949GB encoding *Ssp*DnaB intein. Selectable marker fragments were amplified from plasmids containing these markers. Plasmids created in this study are listed in Supplementary Table 1 with links to Addgene for plasmid sharing and GenBank sequence files.

### HEK293T and U2OS Cell Culture

All cells were cultivated in Dulbecco’s modified Eagle’s medium (DMEM) (Sigma) with 10% fetal bovine serum (FBS)(Lonza), 4% Glutamax (Gibco), 1% Sodium Pyruvate (Gibco) and penicillin-streptomycin (Gibco). Incubator conditions were 37°C and 5% CO2.

### Virus Production

A viral packaging mix of pLP1, pLP2, and VSV-G were co-transfected with each lentiviral vector into Lenti-X 293T cells (ClonTech), seeded the day before in 6-well plates at a concentration of 1.2×10^6^ cells per well, using Lipofectamine 3000. Media was changed 6h after transfection then incubated overnight. 28 hour post transfection, the media supernatant containing virus was filtered using 45 μM PES filters then stored at −80C until use.

### Transduction

The day prior to transduction, U2OS cells were seeded into 12-well plates at a density of 1.5×10^5^ cells per well. Prior to transduction, media was changed to media containing 10 μg/mL polybrene, 1 mL per well. 250 μL of each respective virus (500 μL total for experimental samples with two viruses added) was added to each well and incubated overnight. Media was changed 24-hour post-transduction. Four days post-transduction, cells were split into duplicate plates. Five days post-transduction, media with antibiotics (130μg/mL Hygromycin, 2μg/mL Puromycin, 700μg/mL G418, or 6μg/mL Blasticidin) was added to each respective well of one replicate plate (the other remained under no selection). Antibiotics selection continued for 2 weeks before analysis with flow cytometry.

### Flow Cytometry

Cells were trypsinized, suspended in media then analyzed on a LSRFortessa X-20 (BD Bioscience) flow cytometer using FACSDiVa software, version 8, on an HP Z230 workstation. Fifty thousand events were collected each run.

### Human iPSC culture and nucleofection

Feeder-free KOLF2-C1 were maintained on plate coated with Synthemax ll-SC Substrate (Corning) in StemFlex media (Thermo Fisher Scientific). Subculture was carried out every 4-6 days via Accutase (STEMCELL) detachment method. After plating, 1X RevitaCell supplement (Gibco Life Technologies) was added for 1d to increase cell viability.

For Cas9/gRNA ribonucleoprotein (Cas9/gRNA RNPs) and donor plasmids nucleofection, 4D-Nucleofector™ System (Lonza) was used with the P3 Primary Cell 4D-Nucleofector kit (Lonza). Cells were at 60%-70% confluency at the time of nucleofection. To assemble Cas9/gRNA RNPs, synthetic single guide RNA (Synthego) was resuspended in TE buffer (Synthego) at 2µg/µl, and 8µl of stock solution was mixed with 20µg Cas9 protein before nucleofection. For each reaction, 2 × 10^6^ cells were collected, resuspended in 100µl complete P3 solution and mixed with pre-assembled Cas9/gRNA RNPs as well as donor plasmids DNA. Doxycycline (Sigma) was added 6d after nucleofection at 5µg/mL. 2d after doxycycline was added, 4µg/mL Blasticidin (Sigma) was applied to select cells with resistance. Surviving single colonies were picked and expanded into Matrigel-coated 24-well plate. If surviving colonies were too large to be manually picked as single colony, cells were replated onto new plate at the density of 2,500 cells per 10cm^2^ plate. Blasticidin treatment continued during single colony expansion in 24-well plates.

For genotyping, genomic DNA was extracted using DNeasy Blood & Tissue Kit (QIAGEN), and PCR was performed using the following primers to identify correctly targeted AAVS1 insertions: (i) EGFP-AAVS (fwd: GCCCGACAACCACTACCTGA, rev: GTGAGTTTGCCAAGCAGTCA), (ii) mScarlet-AAVS (fwd: CTGAGGTCAAGACCACCTACAAG, rev: GTGAGTTTGCCAAGCAGTCA).

---

**Supplementary Table 1.**
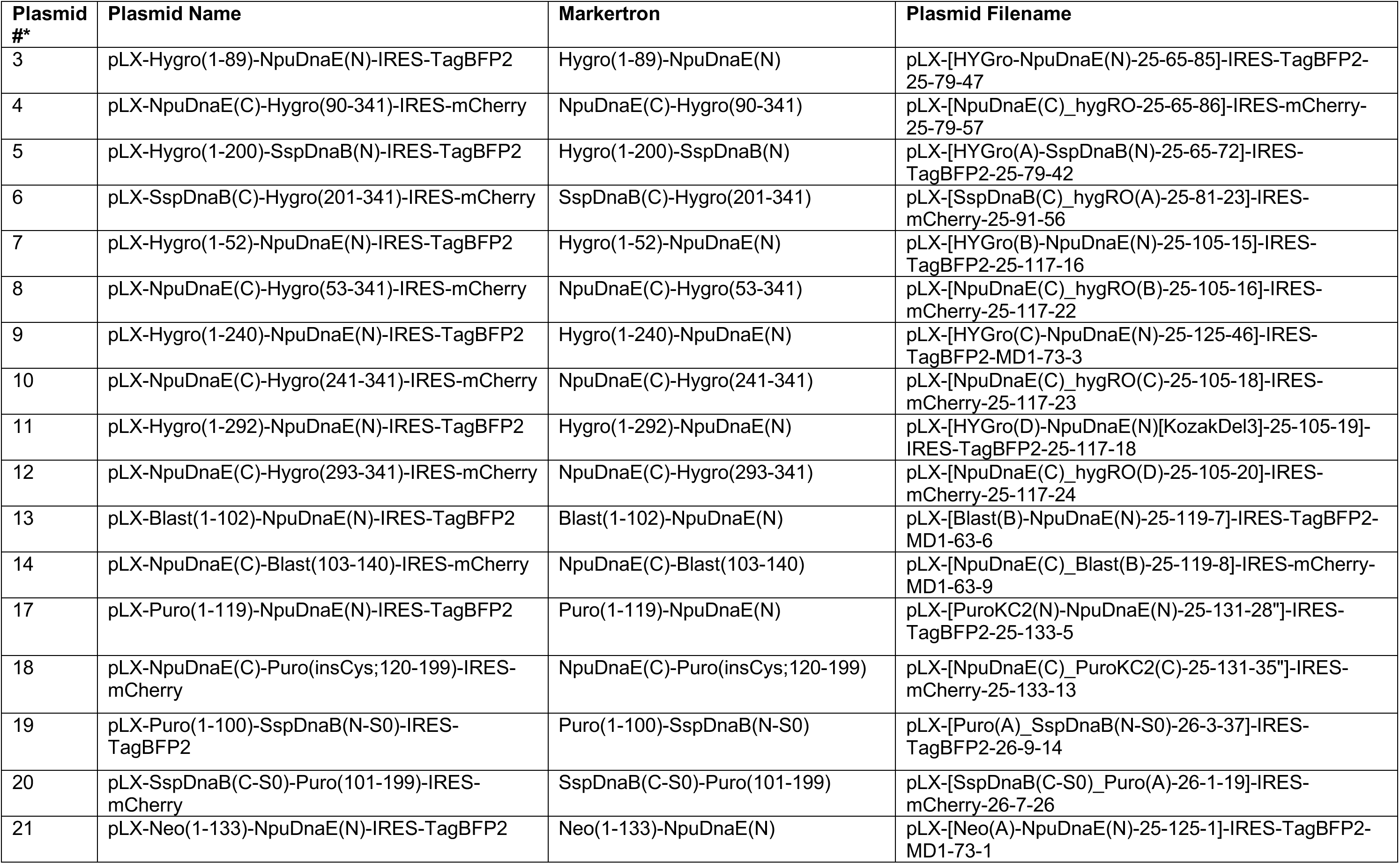
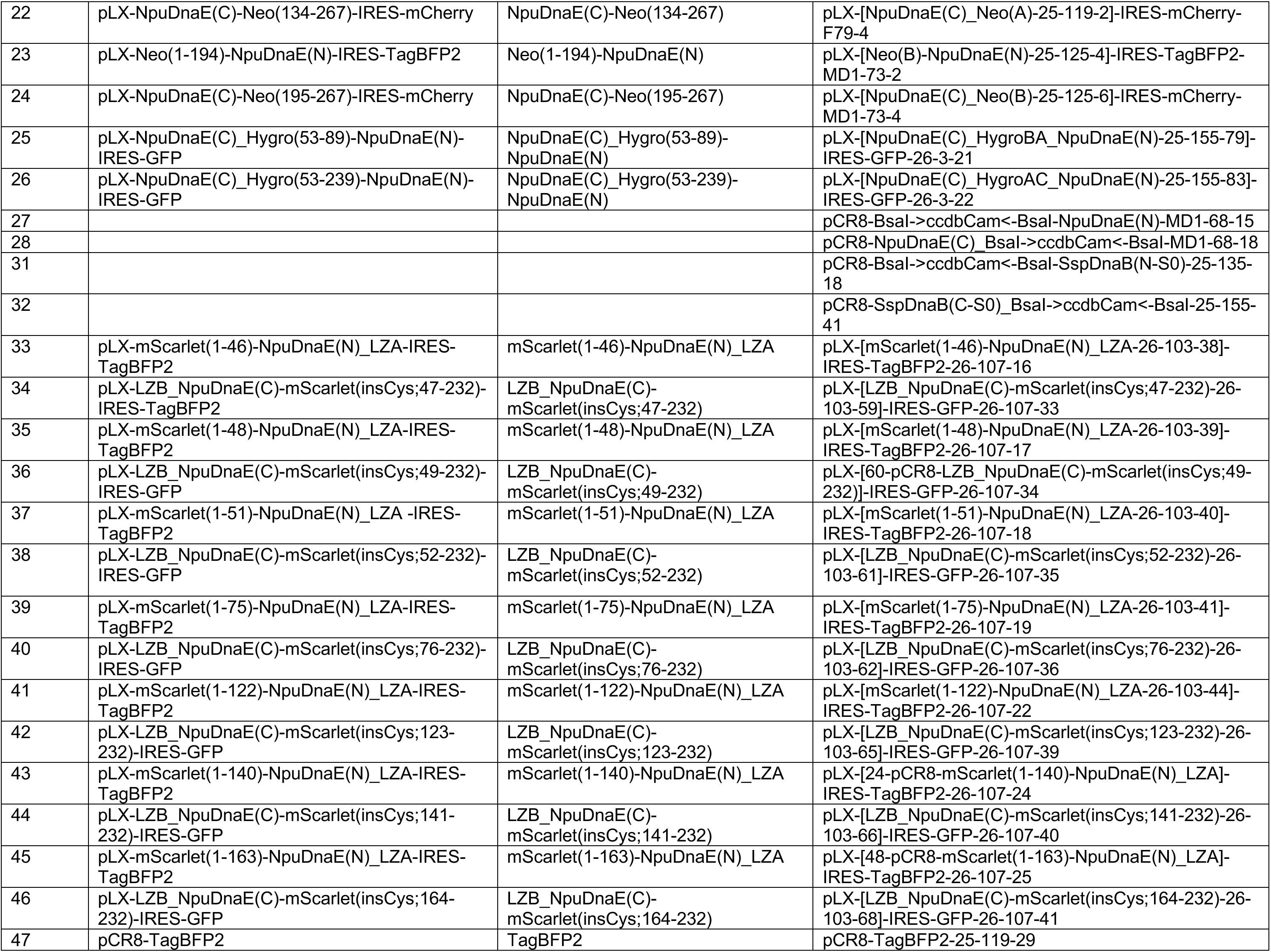
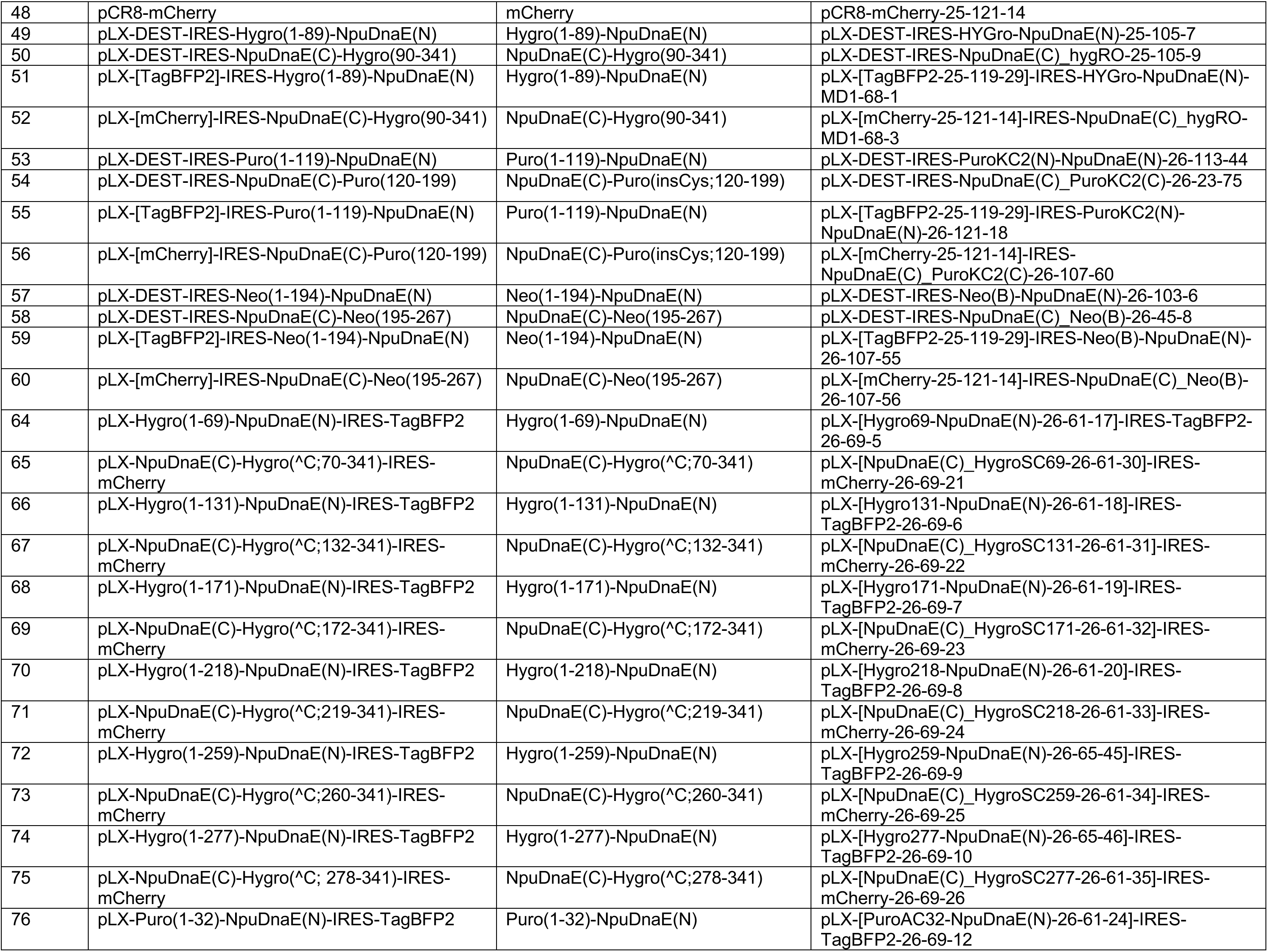
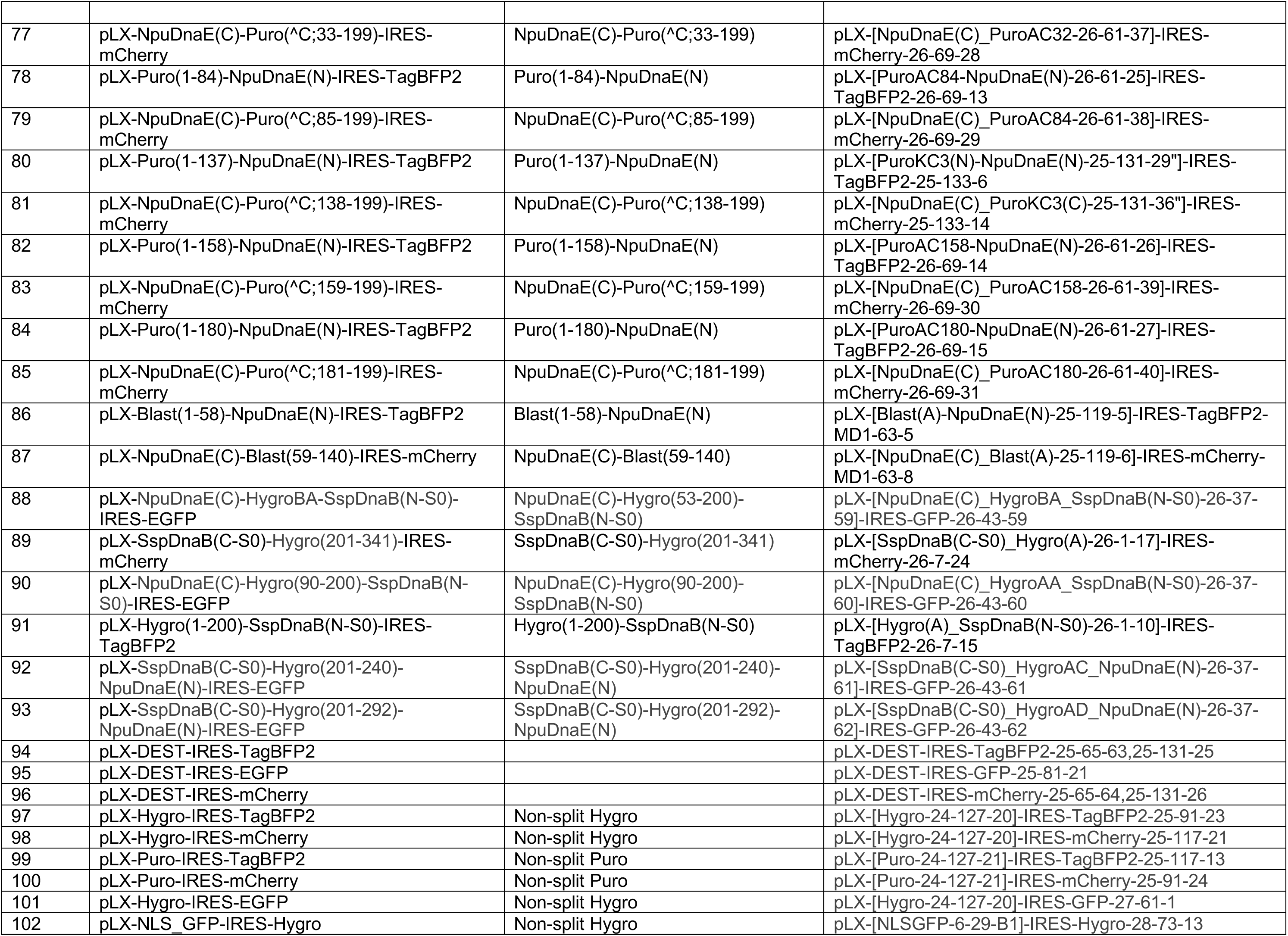
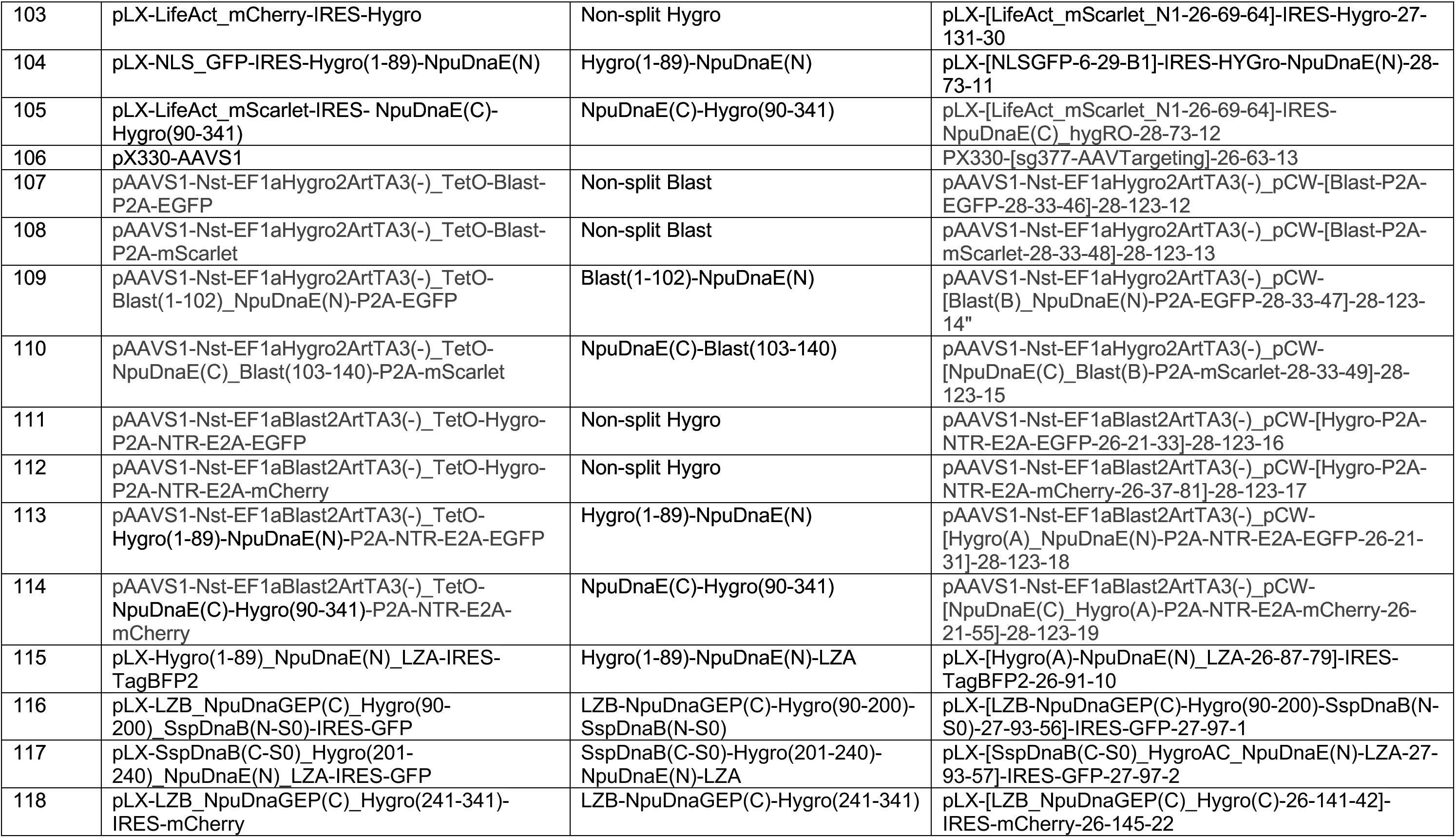
Plasmids will be available on Addgene. For more information: http://intr.es

